# RefKA: A fast and efficient long-read genome assembly approach for large and complex genomes

**DOI:** 10.1101/2020.04.17.035287

**Authors:** Yuxuan Yuan, Philipp E. Bayer, Robyn Anderson, HueyTyng Lee, Chon-Kit Kenneth Chan, Ruolan Zhao, Jacqueline Batley, David Edwards

**Author notes:** These authors contributed equally to this work (Yuxuan Yuan, Philipp E. Bayer, Robyn Anderson).

## Abstract

Recent advances in long-read sequencing have the potential to produce more complete genome assemblies using sequence reads which can span repetitive regions. However, overlap based assembly methods routinely used for this data require significant computing time and resources. Here, we have developed RefKA, a reference-based approach for long read genome assembly. This approach relies on breaking up a closely related reference genome into bins, aligning *k*-mers unique to each bin with PacBio reads, and then assembling each bin in parallel followed by a final bin-stitching step. During benchmarking, we assembled the wheat Chinese Spring (CS) genome using publicly available PacBio reads in parallel in 168 wall hours on a 250 CPU system. The maximum RAM used was 300 Gb and the computing time was 42,000 CPU hours. The approach opens applications for the assembly of other large and complex genomes with much-reduced computing requirements. The RefKA pipeline is available at https://github.com/AppliedBioinformatics/RefKA

## Introduction

A high-quality genome assembly can facilitate a better understanding of genetic variation and gene function in a species (Yuan, et al., 2017). In the past decade, most assemblies generated were fragmented and collapsed in repeat regions due to the short sequence reads used, and consequently, these less accurate genome assemblies have limited our understanding of biological functions encoded in genomes. With the development of long-read sequencing, genome assembly quality is improving with many repetitive genome regions being assembled (Chaney, et al., 2016; Goodwin, et al., 2016; Yuan, et al., 2017). Currently, two real-time long read sequencing technologies are commonly used, manufactured by Pacific Biosciences (PacBio) and Oxford Nanopore Technologies (ONT). The average read length of PacBio long reads and ONT long reads is around 10 Kb to 100 Kb (Chaney, et al., 2016), much longer than short reads (<300 bp) produced by the Illumina sequencing platforms. However, the read accuracy of PacBio and ONT reads is usually much lower than Illumina reads.

Compared to short reads, long reads are better for assembling repetitive and complex genomic regions, with assemblies usually closer to the flow-cytometry estimated genome sizes (Belser, et al., 2018; Hatakeyama, et al., 2017). However, due to the length and high error rate of long reads, their *de novo* assembly is complex and computationally demanding (Ruan and Li, 2019), limiting the uptake of long read sequencing, especially with large and complex genomes where long reads offer the greatest potential.

Different methods have been developed to improve the process of long read *de novo* assembly (Chin and Khalak, 2019; Chin, et al., 2016; Koren, et al., 2017; Lin, et al., 2016; Ruan and Li, 2019; Vaser and Šikić, 2019; Xiao, et al., 2017), though most are still difficult to apply to large and complex genomes. For instance, a recent study of a wheat genome assembly using FALCON took 17 CPU years with a peak memory usage of 1.2 Tb (Zimin, et al., 2017). A recent long read assembler wtdbg2 (Ruan and Li, 2019) reduces computing requirements, though the assembly contiguity is smaller than commonly used assemblers such as Canu (Geest, 2019; Koren, et al., 2017). Flye uses an alternative approach to speed up genome assembly (Kolmogorov, et al., 2019; Lin, et al., 2016), however the large computing memory requirement limits its application. Moreover, a recent benchmark of wtdbg2 and Flye with a plant genome shows that these assemblers produced more misassemblies compared to Canu (Geest, 2019), and so may need further correction prior to downstream analyses.

Here we describe a reference-based approach, RefKA, to reduce the computational requirements, by using unique *k*-mers, string graph and a tiling path approach. The assembly quality from RefKA is comparable with the assemblies produced by state-of-the-art assemblers, such as FALCON, with reduced time and computing requirements.

To demonstrate RefKA, we used public PacBio reads of the bread wheat (*Triticum aestivum*) genome to benchmark the tool and assess assembly quality. The results demonstrate that RefKA is a fast method for assembling large and complex genomes. The success of genome assembly for wheat opens applications to assemble other large and complex reference genomes.

## Results

We have developed RefKA to accelerate long read genome assembly using reduced computational resources. This approach works by breaking a related reference genome into 500 Kb bins with 100 Kb overlaps, then using *k*-mers (k=41) unique for each bin to identify long sequence reads for bin assembly. Each bin is assembled separately in parallel, followed by a final stitching step to produce pseudo-molecules. We evaluated RefKA using a public wheat Chinese Spring (CS) PacBio dataset.

### Allocation of the reference assembly into overlapping 500 Kbp bins

For the assessment, the reference wheat genome (International Wheat Genome Sequencing, et al., 2018) was split into overlapping bins using a 500 Kb sliding window and a 400 Kb step. The number of bins extracted from each pseudo-chromosome is shown in **Figure 1**.

**Figure 1.**
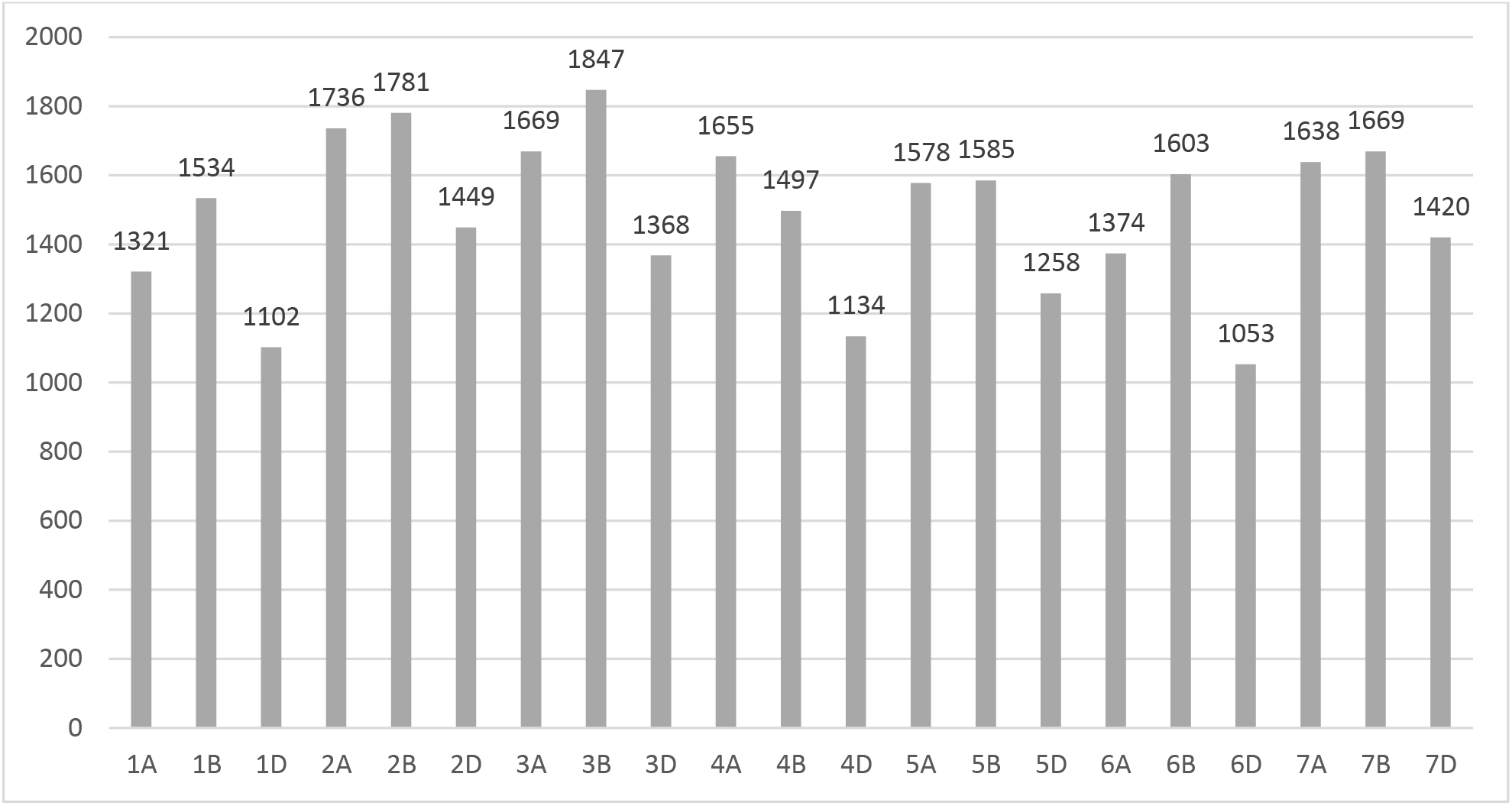
Number of bins extracted from the wheat reference genome assembly (International Wheat Genome Sequencing, et al., 2018) using a 500 Kb sliding window and a 400 Kb step.

### Reference based wheat PacBio genome assembly

After removing PacBio sequence reads shorter than 5 Kbp, 35,265,607 reads with a total length of 496,759 Mbp (~30X) were split into bins and used to reassemble the genome. The resulting assembly consists of 62,157 contigs with an N50 of 415,390 bp (**Table 1**) and a total length of 12,831 Mbp. In comparison, the published FALCON Trit 1.0 assembly (Zimin, et al., 2017) has 35,652 more contigs and around half of the assembly contiguity (N50) (**Table 1**). However, the total assembly size (12,831 Mbp) is slightly smaller than the FALCON Trit 1.0 assembly (12,939 Mbp).

**Table 1.** Comparison of assembly statistics between RefKA and the public FALCON Trit 1.0 (Zimin, et al., 2017) wheat Chinese Spring assemblies.

| Assembly | Number of contigs | Total size without Ns (bp) | N50 (bp) |
| --- | --- | --- | --- |
| **FALCON Trit 1.0** | 97,809 | 12,939,100,857 | 215,314 |
| **RefKA** | 62,157 | 12,831,206,566 | 415,390 |

**Table 2.** Comparison of computational resource used for constructing the RefKA assembly and the FALCON Trit 1.0 assembly (Zimin, et al., 2017).

| Assembly | Contig anchored | CPU hours | Runtime |
| --- | --- | --- | --- |
| **FALCON Trit 1.0** | No | 150,000 | 3 weeks |
| **RefKA** | Yes | 42,000 | 1 week |

### Computational resource consumption

Our approach consumed fewer computational resources than used in the FALCON Trit 1.0 assembly (Zimin, et al., 2017). In addition, the contigs from our assembly are placed within pseudo-chromosomes based on the reference assembly.

### Accuracy of the RefKA pipeline

To demonstrate the accuracy of RefKA, we aligned our assembly with the recently published IWGSC_RefSeqv1.0 assembly (International Wheat Genome Sequencing, et al., 2018) (**Figure 2**; Supplementary Figure 1), demonstrating that the long read assembly produced by RefKA is consistent with the published reference short read assembly.

**Figure 2.**
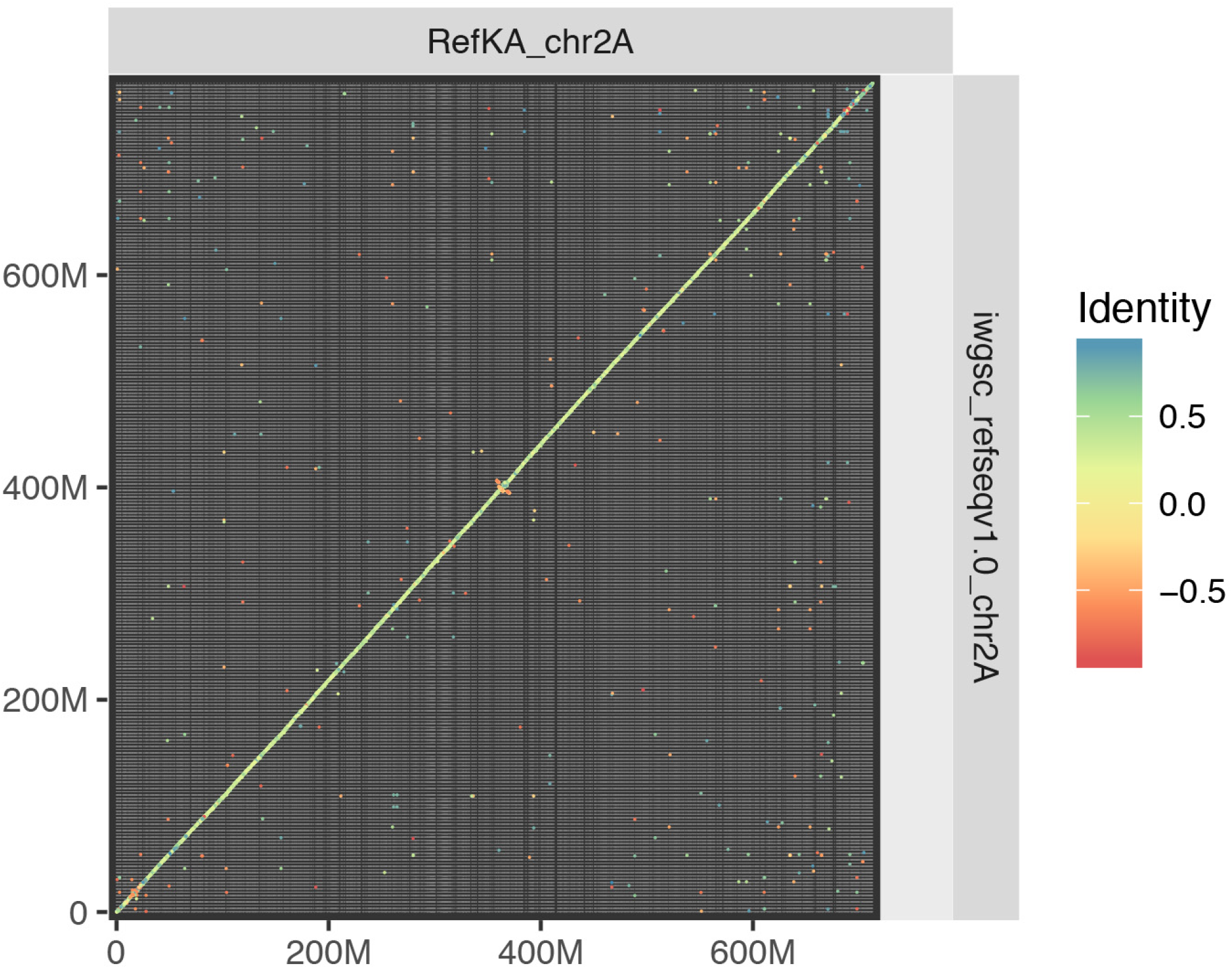
Comparison between the RefKA and IWGSC_refSeqv1.0 assemblies for chromosome 2A.

We polished the RefKA assembly using publicly available Illumina reads and five rounds of ntEdit (Warren, et al., 2019). LTR assembly index (LAI) scores measure how well repetitive regions assemble (Ou, et al., 2018). We compared LAI scores between the unpolished, polished RefKA and IWGSC assemblies. All assemblies had comparable LAI scores between the A, B, and D subgenomes with the RefKA LAI scores being higher in the A subgenomes and slightly higher in the B subgenomes after polishing (**Table 3**).

**Table 3:**
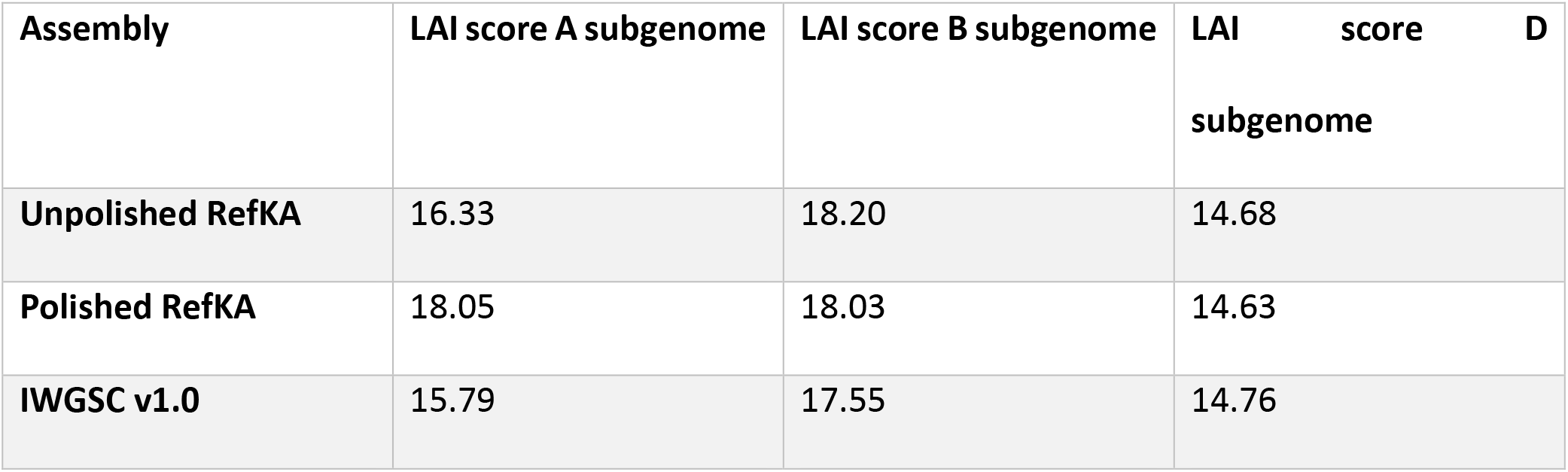
Comparison of LAI scores between unpolished, polished RefKA and IWGSC assemblies per subgenome

| Assembly | LAI score A subgenome | LAI score B subgenome | LAI score D subgenome |
| --- | --- | --- | --- |
| Unpolished RefKA | 16.33 | 18.20 | 14.68 |
| Polished RefKA | 18.05 | 18.03 | 14.63 |
| IWGSC v1.0 | 15.79 | 17.55 | 14.76 |

## Discussion

With advances in data quality and reducing cost, long read sequencing has been increasingly used in genomic studies. However, due to the high error rates and long read length, the routinely used overlap methods struggle to manage the substantial quantity of data required to assemble large and complex genomes without significant computational resources. Different approaches, such as the fuzzy*-Bruijn* graph (Ruan and Li, 2019) and repeat graph (Kolmogorov, et al., 2019), are being applied to address this problem, however the assembly quality is lower than mainstream methods such as Canu and FALCON (Geest, 2019). To address this problem and to make use of the increasing number of available reference assemblies, we have developed a reference-based genome assembly approach for large and complex genomes.

*K*-mer size selection is important for RefKA. A large *k*-mer size has the potential to reduce the possibility of alignment due to the high error rate in long reads, while a small *k*-mer size will decrease *k*-mer specificity leading to few unique *k*-mers. For the public wheat PacBio dataset, we used k=41 as a trade-off between accuracy and false positive alignments. The optimal *k*-mer value will differ between datasets and genomes and is likely to increase as the error rate of long read sequencing continues to decrease, as well as decrease when assembling genomes which are more distant from the reference genome. Due to the errors in long sequence reads, some reads may be allocated to the incorrect bin, though with the minimum read coverage requirements of Canu, it is unlikely that these reads would contribute to the bin assembly, and so we take a redundant approach where reads may be allocated to more than one bin.

Currently, as RefKA is based on a reference genome to define bins, it cannot identify large scale structural variation between the assembled and the reference genome. However, further data such as produced by HiC (Lieberman-Aiden, et al., 2009) and optical mapping (Schwartz, et al., 1993) may help to correct and validate such variation. Furthermore, large regions which are absent from the reference genome will not be assembled. However, it may be possible to assemble these separately after collating reads which do not map to the assembly.

Comparative benchmarking demonstrates that our approach can reduce the computational resources used for genome assembly using long reads compared to the strict *de novo* approach, with higher contiguity of assembled contigs. LTR Assembly Index (LAI) were 18, 18, and 14 for the three subgenomes in the RefKA assembly indicating ‘reference’ quality of the assembly (Ou, et al., 2018), which is comparable to the IWGSC assembly. Interestingly, polishing greatly improved the A subgenome LAI scores while it slightly reduced the B subgenome LAI score. The slightly smaller assembly size in this example may be due to the strict assembly parameters applied. The unique *k*-mer basis of this reference-based approach can be used for the development of a *de novo* assembly method. Following the identification of unique *k*-mers from either PacBio reads or a second Illumina dataset, a graph can be generated based on reads sharing these *k*-mers, with subsequent binning and assembly.

## Conclusions

To make use of the increasing number of reference genome assemblies and the reducing cost of long read sequence data, we have developed an approach to produce genome assemblies using less computational resources compared to current *de novo* assembly methods. The application of RefKA for the assembly of the large and complex bread wheat genome demonstrates the use of this approach which can be applied to additional datasets as they become available.

## Acknowledgements

We acknowledge the supercomputing resources provided by the Pawsey Supercomputing Centre with funding from the Australian Government and the Government of Western Australia. This work was supported by the Australian Research Council Grant Nos. DP160104497, LP160100030, LP140100537.

## Author contributions

D.E conceived and directed the research. All authors contributed to the development of the tool. P.B., K.C and Y.Y coded the tool. Y.Y. wrote the manuscript with contributions from other authors. RZ carried out polishing and quality assessment of the assemblies. All authors read and approved this manuscript.

## Methods

### Data and code availability

RefKA is available at https://github.com/AppliedBioinformatics/RefKA. RefKA requires Jellyfish (Marcais and Kingsford, 2011), SOAP2 (Li, et al., 2009), SOAP3 (Liu, et al., 2012) and Canu (Koren, et al., 2017).

The PacBio dataset tested is available at NCBI Sequence Read Archive (SRA) database SRR5816161 under project PRJNA392179.

### Methods implemented in RefKA

RefKA is designed to assemble a large and complex genome in a fast and efficient way using long-read sequencing data by identifying unique *k*-mers in a related reference genome. RefKA implements the following workflow (**Figure 3**):

1. RefKA identifies *k*-mers that are unique in a related reference genome and numbers the *k*-mers. The optimal *k*-mer size is selected by testing a range of *Ks* using a small proportion of the raw reads.
2. After unique *k*-mer identification, RefKA aligns the *k*-mers to the reference genome. *K*-mers with perfect alignment (no gaps, no mismatch and only one alignment position) are considered and assigned with coordinates. Using a 500 kb sliding window and a 400 Kb steps on the reference genome, the anchored *k*-mers are grouped into different bins.
3. RefKA aligns the *k*-mers allocated in each bin to the long-read sequencing dataset. Due to the limitations of maximum number of nucleotides in an index, the entire long-read dataset is chunked by RefKA. The mapping process is repeated for each chunk.
4. Long sequence reads are extracted and allocated into different bins using the anchor information of each unique *k*-mer on the reference genome. During allocation, if a read can be aligned by multiple *k*-mers from different chromosomes, the read is assigned into the chromosome with the higher number of corresponding *k*-mers.
5. RefKA *de novo* assembles the long reads allocated in each bin. During assembly, read correction and trimming are performed.
6. RefKA uses the known *k*-mer anchor information to stitch contigs in each chromosome. This step connects adjacent bins. Since adjacent bins overlaps by 100Kb, it is possible to re-align unique *k*-mers with contigs in adjacent bins, and then merging them automatically. Overlapping contigs are merged based on simple overlaps, with the longer contig taking precedence.

**Figure 3.**
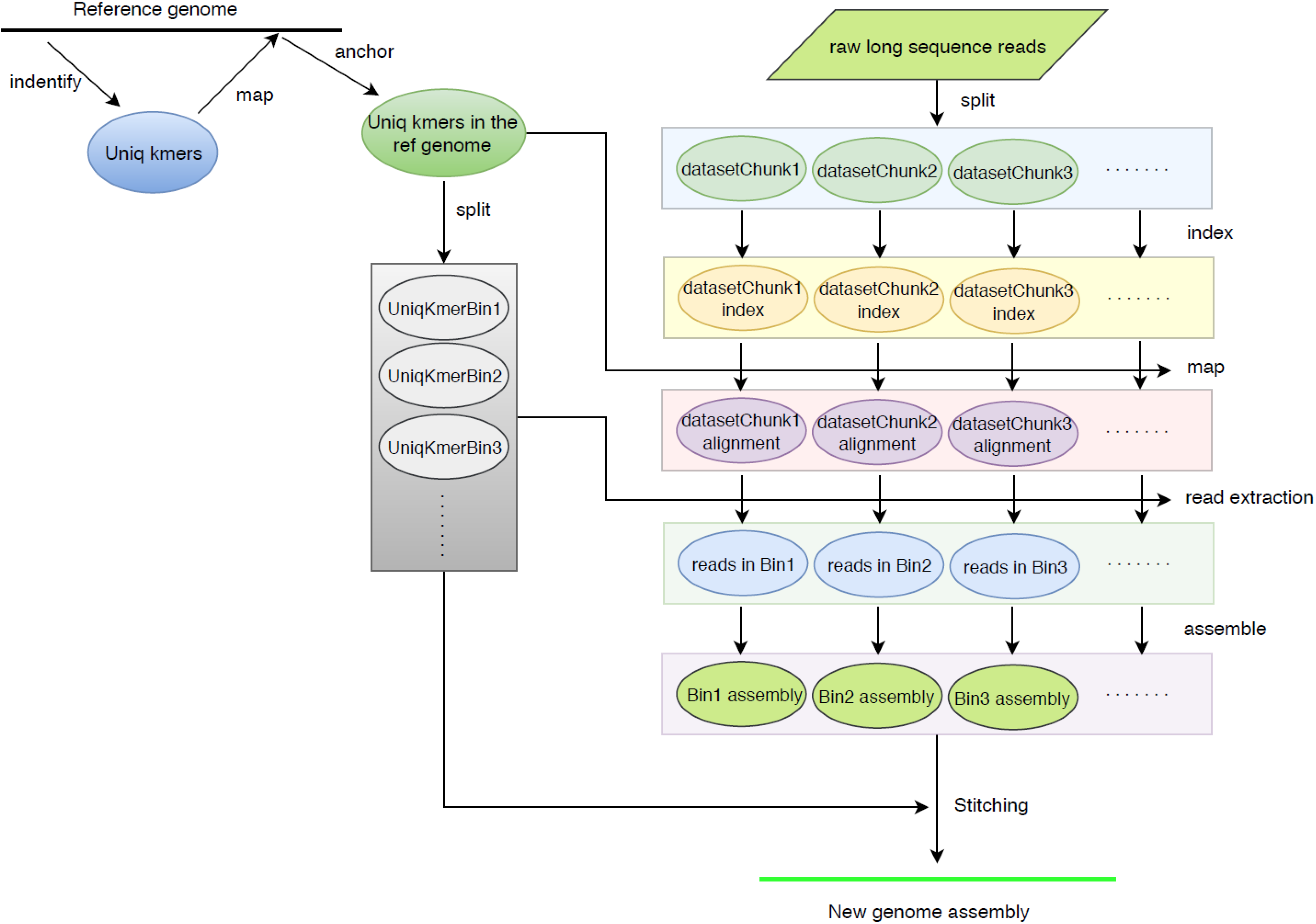
the workflow used in the *k*-mer based genome assembly approach.

### Quality assessment

The RefKA genome was improved using five rounds of polishing using publicly available wheat data for Chinese Spring using ntEdit (Warren, et al., 2019) using data from (Zimin, et al., 2017) (BioProject ID: PRJNA392179). The quality of repeat assembly was assessed using LTR_retriever v2.8 (Ou, et al., 2018).

### Benchmarking

During benchmarking, RefKA was used to assemble the genome of bread wheat cultivar Chinese Spring. The reference genome used was the one recently released by the International Wheat Genome Sequencing Consortium (IWGSC) (International Wheat Genome Sequencing, et al., 2018). The raw PacBio reads were downloaded from SRA (SRR5816161) (Zimin, et al., 2017). The read coverage is around 36X. Computational resources were provided by the Pawsey Supercomputing Centre. For array jobs, RefKA was tested on Magnus cluster with two Intel Xeon E5-2690 v3 12-core CPUs per node. For normal jobs, RefKA was tested on Zeus cluster with Intel Xeon E5-2680 v4 28 cores per node.

During unique *k*-mer identification, 41 was selected as the *k*-mer size after an initial test using different *k*-mer sizes. PacBio reads that are less than 5 Kb long were removed from the original dataset to ensure that there are enough unique *k*-mers can map to the reads. During assembly, the computing resources used, CPUs, RAM, and run time were collated. After stitching, minimap (Li, 2016) and minidot (https://github.com/thackl/minidot) (‘-L 5000 -c 30 -f 0.01’) were used to align the assembled contigs to the reference genome and check the accuracy of RefKA assembly.

